# Identification of genetic drivers of plasma lipoproteins in the Diversity Outbred mouse population

**DOI:** 10.1101/2023.08.26.554969

**Authors:** Tara R. Price, Christopher H. Emfinger, Kathryn L. Schueler, Sarah King, Rebekah Nicholson, Tim Beck, Brian S. Yandell, Scott A. Summers, William L. Holland, Ronald M. Krauss, Mark P. Keller, Alan D. Attie

## Abstract

Despite great progress in understanding lipoprotein physiology, there is still much to be learned about the genetic drivers of lipoprotein abundance, composition, and function. We used ion mobility spectrometry to survey 16 plasma lipoprotein subfractions in 500 Diversity Outbred (DO) mice maintained on a Western-style diet. We identified 21 quantitative trait loci (QTL) affecting lipoprotein abundance. To refine the QTL and link them to disease risk in humans, we asked if the human homologues of genes located at each QTL were associated with lipid traits in human genome-wide association studies (GWAS). Integration of mouse QTL with human GWAS yielded candidate gene drivers for 18 of the 21 QTL. This approach enabled us to nominate the gene encoding the neutral ceramidase, *Asah2*, as a novel candidate driver at a QTL on chromosome 19 for large HDL particles (HDL-2b). To experimentally validate *Asah2*, we surveyed lipoproteins in *Asah2^-/-^*mice. Compared to wild-type mice, female *Asah2*^-/-^ mice showed an increase in several lipoproteins, including HDL. Our results provide insights into the genetic regulation of circulating lipoproteins, as well as mechanisms by which lipoprotein subfractions may affect cardiovascular disease risk in humans.

## Introduction

High concentrations of LDL-cholesterol (LDL-C) are associated with increased cardiovascular disease (CVD) risk. Interventions to reduce LDL-C result in improved cardiovascular outcomes. Small dense LDL particles are associated with CVD, including coronary artery disease and stroke (1), and are strong predictors of cardiovascular events (2). Likewise, larger HDL-2b particles are better predictors of coronary heart disease than small, dense HDL-3, LDL-C, or HDL-C levels (3–5). Variation in protein and lipid composition, and particle size can affect lipoprotein function (6–9).

Genetically diverse mouse populations, resulting from intercrossing or outcrossing inbred strains, can be used to discover the genetic drivers of lipoprotein abundance and composition. For example, intercross mouse populations identified gene loci affecting apolipoprotein (Apo) A2 (10). In an intercross study involving the RIIIS/J and 129S1/SvImJ mouse strains, eight unique loci affecting plasma cholesterol and causative genes within the loci were identified (11). Panels of recombinant inbred mouse strains have been used to leverage naturally occurring polymorphisms, which showed heterogeneity in lipoprotein size and apolipoprotein composition (12). The hybrid mouse diversity panel (HMDP), a collection of 100 mouse strains, utilizes natural strain variation in a systems genetics approach to identify genetic drivers of phenotypes (13). A meta-analysis of nearly 5000 HMDP mice identified 26 significant loci associated with HDL-C. Several loci, including one for ApoA2, were consistent with previous reports, while other loci provided novel insights into gene-environment interactions (14).

The use of outcrossed mouse populations, including the Collaborative Cross (CC), brings additional genetic diversity to mouse genetic screens. In a study using 25 CC strains, increased adiposity and liver steatosis were associated with increasing total, HDL, and LDL cholesterol (15). Key genetic regulators of hepatic lipids were linked to diet-induced changes in liver steatosis severity and plasma lipid measures. Leveraging the genetic diversity of the Diversity Outbred (DO) mouse population, an outbred stock derived from eight founder strains of the CC, three loci were identified for plasma cholesterol (16).

In the present study, we utilized the DO mouse population to determine quantitative lipoprotein subclasses, identify quantitative trait loci (QTL) for several subclasses, and nominate candidate genetic drivers of lipoprotein particle sizes. We identified several known cholesterol-related genes, *Apoa2* and *Foxo1*, as well as novel loci associated with plasma cholesterol.

## Methods

### Animal husbandry – Founder and DO mice

All animal protocols were approved by the Animal Care and Use Committee at University of Wisconsin-Madison. The eight founder strains (C57BL/6J (B6); A/J; 129S1/SvImJ (129); NOD/ShiLtJ (NOD); NZO/HILtJ (NZO); PWK/PhJ (PWK); WSB/EiJ (WSB); and CAST/EiJ (CAST)) and DO mice were purchased from Jackson Laboratories (Bar Harbor, ME) and maintained at the University of Wisconsin-Madison, as previously described (17, 18). Briefly, founder mouse strains were fed standard laboratory chow (Formulab Diet 5008, LabDiets, Brentwood, MO) or given a high-fat, high-sucrose diet (TD.08811, Envigo, Madison, WI) for 18 weeks. Diversity Outbred (DO) mice were maintained on the same high-fat, high-sucrose diet as the founder mice for 16 weeks. Animals were euthanized at 22 weeks of age and plasma collected and stored at -80°C.

### Animal husbandry – *Asah2* mice

*Asah2* mice were generated by Richard Proia (NIH)(19) and provided to the Summers/Holland lab. All animal procedures were performed in compliance with protocols approved by the Institutional Animal Care and Use Committee (IACUC) at the University of Utah and adhered to National Institutes of Health (NIH) standards. Male (*n* = 5-6 per genotype) and female (*n* = 4-7 per genotype) mice were maintained under standard laboratory conditions at a temperature of 22°C to 24°C, in groups of 2 to 5 mice, with a 12-hour light/dark cycle. Mice were allowed *ad libitum* access to food and water unless fasting conditions were required for experimental procedures. Animals were fed a normal chow diet from the age of 4 weeks and transitioned to a high-fat diet (60% total energy; D12492; Research Diets Inc, New Brunswick, NJ, USA) at 9 weeks of age for 16 weeks. At 25 weeks of age, mice were anesthetized with isoflurane and blood collected by cutting the brachial artery. Whole blood was collected into vacutainers coated with 20% K_2_EDTA, centrifuged at 7500xg for 7.5 minutes, and plasma separated. Plasma was stored at -80°C.

### Plasma lipoprotein fractionation by ion mobility analysis

To analyze lipoprotein class size, plasma lipoproteins were separated by ion mobility analysis as previously described (20, 21). Briefly, lipoproteins were harvested on paramagnetic particles, washed to remove free salt and proteins (*e.g.*, IgG, albumin, and transferrin), and then resuspended in 25LJmM ammonium acetate. Lipoproteins were then fractionated and quantified by summing the total number of particles within specific size ranges. Supplemental Table S1 shows the lipoprotein subclasses, their size ranges, and nomenclature.

### QTL mapping of plasma lipoprotein phenotypes

Mapping of plasma lipoproteins for quantitative trait loci analysis was performed as previously described (17). Briefly, 478 Diversity Outbred (DO) mice (236 female, 242 male) were obtained from Jackson Laboratories (Bar Harbor, ME) and maintained on a high-fat, high-sucrose diet (TD.08811, Envigo, Madison, WI) for 16 weeks and plasma collected for lipoprotein sizing by ion mobility analysis. Lipoprotein phenotype data were rankz-transformed to achieve a normal distribution prior to mapping. Genetic mapping was performed using the qtl2 package with kinship correction to identify quantitative trait loci (QTL) using the GRCm38 genome build and Ensembl 75 for gene annotation. Genome scans used sex, mouse cohort (wave), and technical batch as additive covariates. As previously described, logarithm of odds (LOD) thresholds were defined through permutation testing to establish a genome-wide family-wide error rate (FWER) for genome-wide QTL (22, 23). A LOD greater than 6.0 was used as the threshold for identifying suggestive QTL and a LOD greater than 7.4 identified significant QTL.

### RT-PCR

For liver gene analyses, liver samples were homogenized in Qiazol lysis buffer in a TissueLyser II and RNA isolated with RNeasy Mini Kit (Qiagen) following manufacturer’s protocols. Hepatic gene expression of *Asah2* was normalized to β-actin (*Actb)* and fold change relative to wild-type controls for each sex were calculated using the 2^−ΔΔCt^ method (24). *Asah2* primer sequences: GATCCATTC TGGGACACTCTTC (Forward), TCCACTGTGAAGCAGGATTG (Reverse). *Actb* primer sequences: AGATGTGGATCAGCAAGCAGG (Forward), TGCGCAAGTTAGGTTTTGTCA (Reverse).

### Statistical Analyses

Founder plasma lipoprotein data were analyzed in JMP Pro version 15.0.0 (SAS Institute, Cary, NC). All data were log-transformed prior to statistical analysis. Chow-fed and HFHS-fed data were initially analyzed separately for strain and sex effects, prior to assessing the interaction effect of diet on strain and sex. Strain, sex, and diet interactions for each lipoprotein subclass was tested using a standard least squares model with p<0.05 denoting a significant effect. Least square means differences with Tukey’s HSD post-hoc analysis determined statistical differences between groups (p<0.05). For instances where the interaction effect and one of the main effects failed to reach significance, a one-way ANOVA was used. Heritability calculations were conducted in R (version 4.3.1) using the “lme4” package to fit a linear mixed model with restricted maximum likelihood. Chow-fed and HFHS-fed mice were analyzed separately, with sex as the fixed effect and strain as the variable effect. *Asah2* mouse data were first checked for a Gaussian distribution and log-transformed if not normally distributed. Statistical analyses of normally distributed data were performed by ANOVA followed by Tukey’s post-hoc analysis. Differences were considered significant at p<0.05.

## Results

### Strain and sex dependence of diet-induced alterations in lipoproteins

To estimate heritability (*h^2^*) in lipoprotein phenotypes in DO mice, we first analyzed the lipoproteins of the eight founder strains of DO mice. The mice were fed a chow or a high-fat, high-sucrose (HFHS) diet and their sera were analyzed by ion mobility analysis, a method that quantitates the various size categories of lipoproteins (**Table S1**) (25).

Mice fed a standard chow diet showed marked strain-dependent differences in their lipoprotein size distribution (**Fig. 1A**, **Fig. S1**), with significant interactions between strain and sex for all lipoprotein subclasses (p<0.03). NOD, NZO, and PWK mice had increased lipoprotein abundance relative to the other strains (p<0.005), with NZO having the highest concentrations of small LDL (LDL-IVa, -IVb, -IVc, p<0.0001). PWK mice showed variation in classes by sex where females (F) had increased the small lipoprotein particles: HDL-3,2a, HDL-2b and midzone relative to male (M) PWK mice (p<0.007, p<0.0001, p<0.0001, respectively). NZO mice had increased concentrations of LDL particles compared to the other strains (p<0.0001), whereas B6, WSB, and CAST mice had the lowest LDL particle concentrations (p<0.03). Heritability was high for all particles, with strain explaining up to 77% of phenotypic variance (**Fig. 1C**, *h^2^* = 0.49 – 0.77).

**Figure 1:**
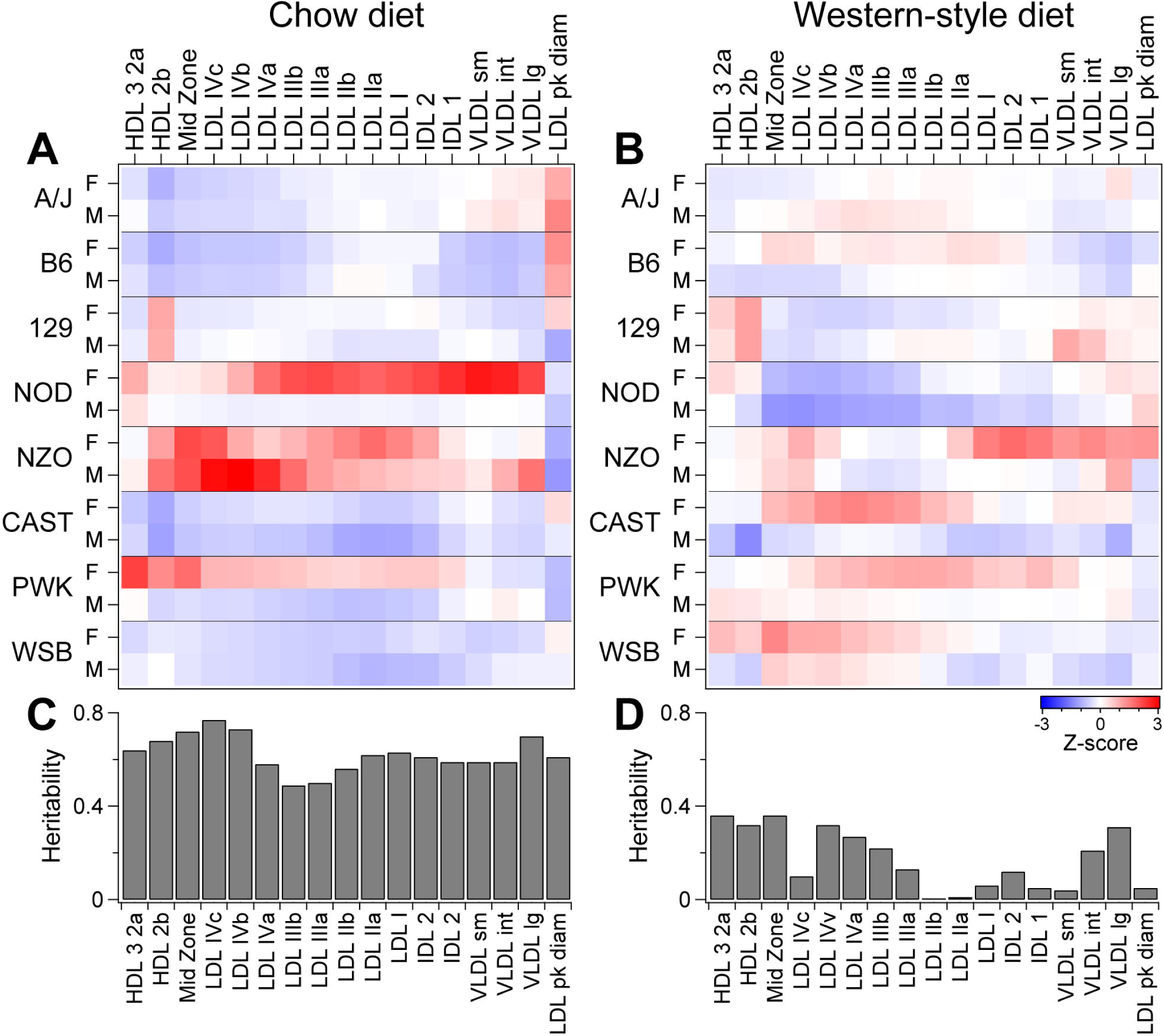
Genetics and diet exert a strong influence on circulating lipoproteins. Profile of lipoprotein subclasses in female (F) and male (M) mice of the eight Diversity Outbred (DO) founder strains maintained on standard rodent chow diet (**A**) or a Western-style diet high in fat and sucrose (**B**). Heritability (*h^2^*) estimates for lipoprotein particles in mice fed a chow diet **(C)** versus a Western-style diet **(D)**.

When placed on the HFHS diet for 16 weeks, total lipoprotein concentrations were increased by ∼50% (25±1 nM) in five of the eight mouse strains, compared to chow-fed mice (17±1 nM, p<0.0001). NZO, NOD, and PWK mice did not significantly change their total lipoprotein particle concentrations in response to the HFHS diet. However, there was no effect of sex or interaction between sex and strain on any of the individual subclasses in response to the HFHS diet (**Fig. 1B**, **Fig. S1**, p>0.3). Therefore, we analyzed the particles by one-way ANOVA to determine differences between strains.

NOD mice had the lowest LDL-IIIb and LDL-IV particle concentrations relative to the other strains (p<0.001). Large LDL-I and LDL-II particles did not vary by strain (p>0.5). HDL-2b concentrations were highest in 129 mice and significantly different from CAST (p<0.0004). Interestingly, the average LDL diameter was higher in chow-fed mice (204±1 Å) compared to HFHS-fed mice (194±1 Å, p<0.0001). LDL peak diameter showed a strong strain dependence for chow-fed mice (p<0.0001) that was not present for HFHS-fed mice (p>0.6). Two mouse strains, A/J and B6, had the largest LDL diameter on the chow diet (p<0.0001), but were not different from other strains on HFHS diet. Heritability for lipoprotein particle concentrations was diminished on the HFHS diet (**Fig. 1D**), with strain explaining ∼40% or less of the variance in particle concentrations (*h^2^* = 0-0.36).

### Genetic association of lipoprotein classes in DO mice

We surveyed all lipoprotein subclasses in ∼500 DO mice genotyped at ∼69,000 genome-wide SNPs, enabling us to identify 30 quantitative trait loci (QTL) with LOD>6.0 (genome-wide p=0.2) for association with plasma lipoprotein subclasses (**Fig. 2A, Table S2**). For several QTL, multiple lipoproteins co-mapped, including one on Chr 10 for LDL-I through LDL-IIIb.

**Figure 2:**
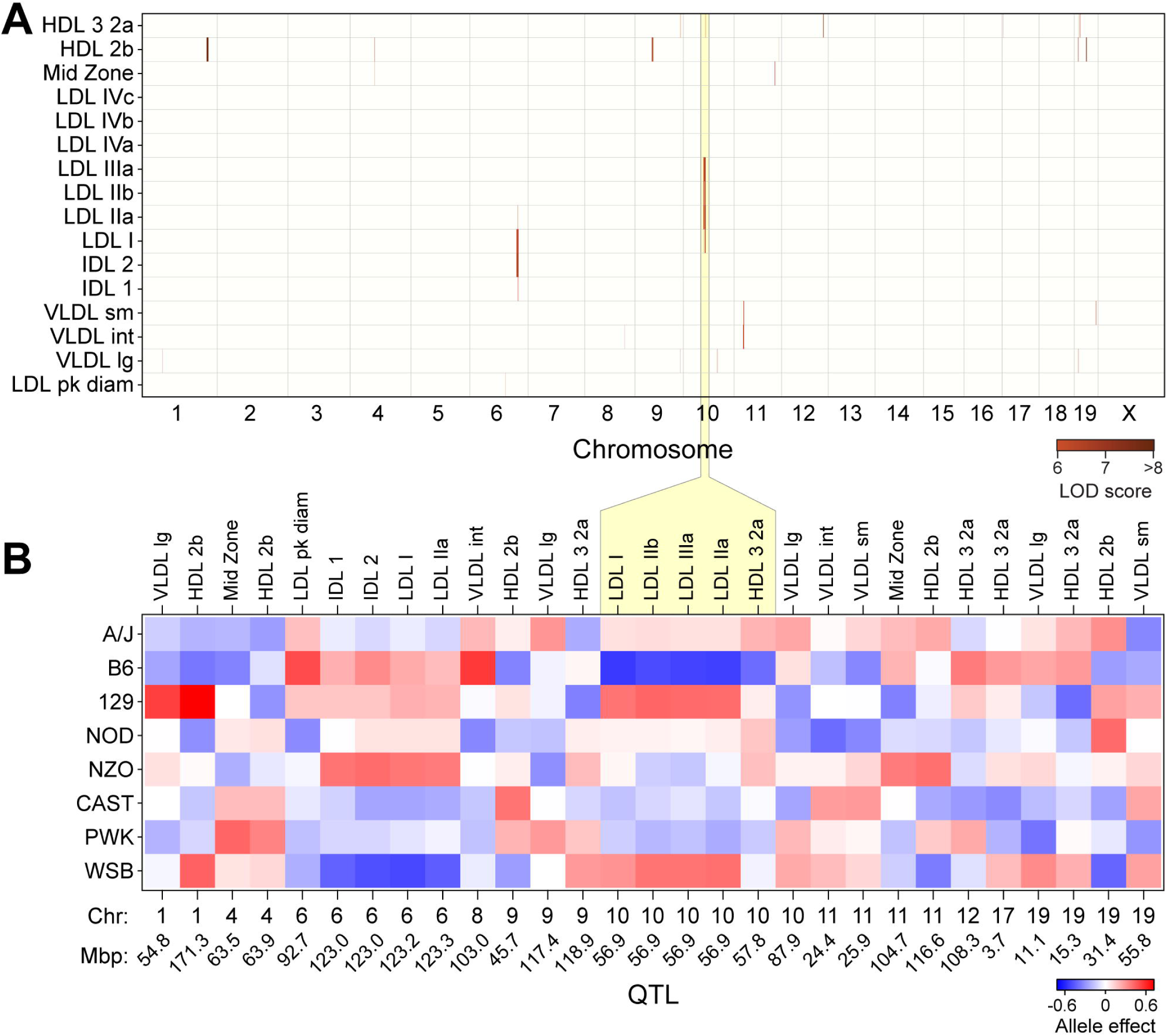
Genetic architecture of circulating lipoproteins. Heatmaps illustrate average Z-scores across all mice for 3-5 mice per sex/strain. Genome-wide QTL for lipoprotein subclasses in ∼500 DO mice maintained on Western-style diet (**A**). Supplemental Table S2 lists all QTL, their genomic positions, LOD scores and allele effect values. Allele effect values are illustrated for 30 QTL for individual lipoprotein subclasses(**B**). Blue depicts alleles associated with reduced lipoprotein values, red for increased values. Founder strains are listed on the left, while lipoproteins and their QTL (Chr and Mbp location) are shown along the top and bottom, respectively. Loci with similar allele effect patterns (*e.g.*, Chr 10 at ∼57 Mpb for LDLIIIb – LDLIIa and HDL-3,2a), are considered one QTL.

Because DO mice derive from an outcross of eight founder strains, there are up to eight alleles segregating among DO mice across the genome. Through haplotype reconstruction, we could determine the association of strain-specific haplotype blocks with each phenotype and thus determine the directionality of each allele’s influence on the phenotype, *i.e.*, their allele signatures. **Fig. 2B** illustrates the allele signatures for all lipoprotein QTL.

We defined lipoprotein QTL based on the genome location of the SNP having the highest LOD score, and the allele signatures of co-mapping traits. For example, a single locus on Chr 10 at ∼57 Mbp that showed five LDL subclasses co-mapping in response to the same allele signature (low for B6 and high for 129 and WSB) was classified as a single QTL. After refining QTL based on genomic proximity and allele effects, we identified 21 unique QTL for all lipoprotein classes. Six QTL were identified for HDL-2b and five QTL for the HDL-2a,3 subclasses, the most for individual subclasses in our analyses (**Table S2**). Large VLDL had four QTL, while the remaining subclasses had one to two QTL.

To nominate causal genes at each QTL, we identified probable genes based on SNP association plots at each locus. One lipoprotein subclass, HDL-2b, significantly mapped to 6 loci, with 4 QTL having a LOD>6.0 (**Fig. 3A**). Each locus had a unique allele signature **(Fig. 3B**), indicating independent genetic regulation of this lipoprotein. By integrating mouse SNPs at each locus (**Fig. S2, Table S3**), candidate genetic drivers were identified. The QTL on Chr 1 was located near *Apoa2*, the second most abundant apolipoprotein component of HDL (26). The QTL on Chr 9 was located near several apolipoproteins, including Apolipoprotein A1 (*Apoa1*), as well as proprotein convertase subtilisin/kexin type 7 (*Pcsk7*). APOA1 is the most abundant apolipoprotein in HDL (26, 27) and has been well characterized for its role in HDL function (28). The association of three genetic variants of *PCSK7* with HDL-cholesterol and acute coronary syndrome have also been recently published (29).

**Figure 3:**
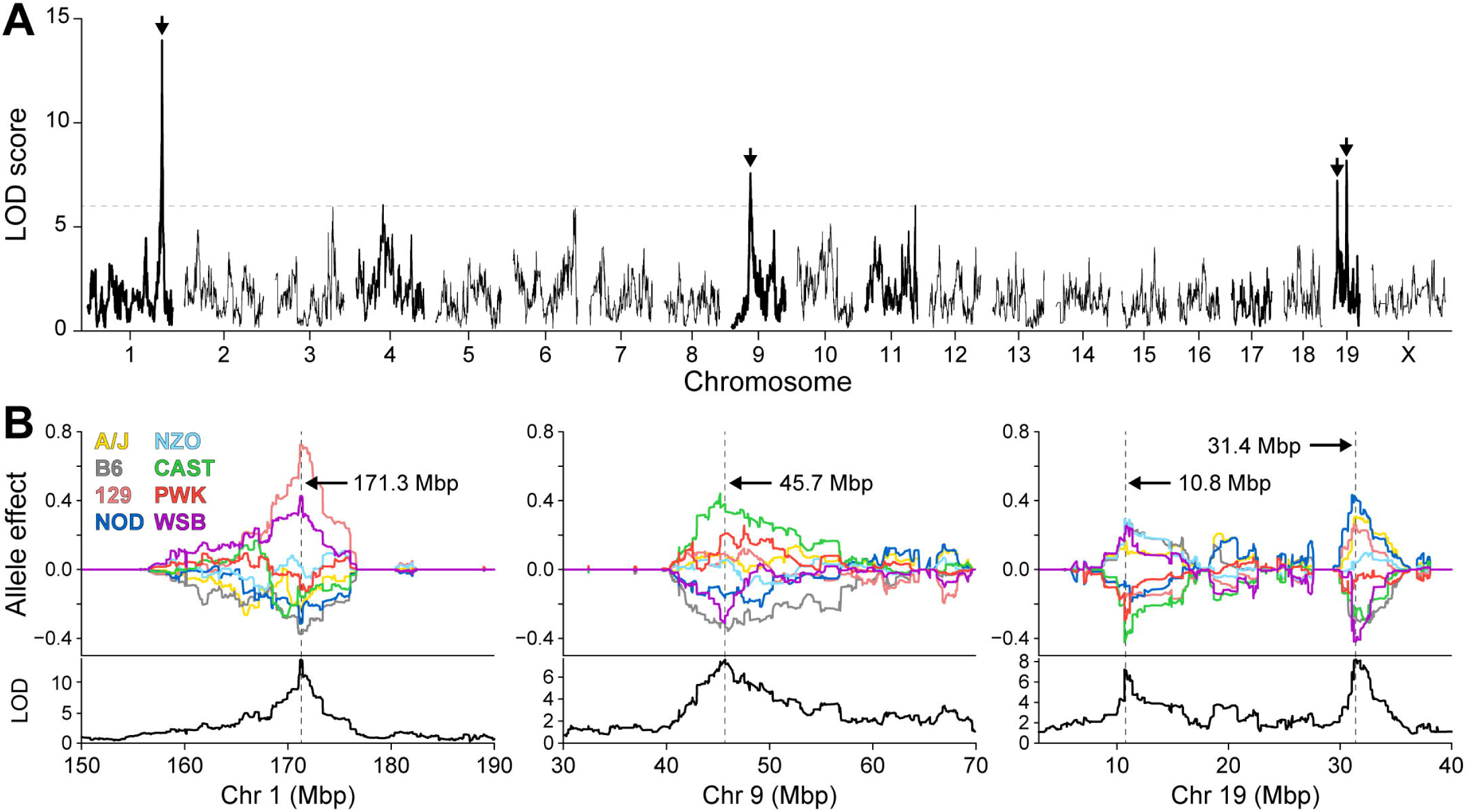
Genetic regulation of circulating HDL-2b. Genome-wide LOD profile for an HDL subclass (HDL-2b) identifies significant QTL on three chromosomes: 1, 9 and 19 (**A**). Allele effect plots illustrate distinct genetic architecture at each locus (**B**). Colored lines represent alleles derived from founder strains. Genomic position for peak SNP listed for each QTL.

Finding QTL at the *Apoa2* and *Apoa1/Pcsk7* loci demonstrated that our genetic screen could identify known drivers of cholesterol and lipoprotein levels. We next turned our attention to two distinct HDL-2b QTL on Chr 19. The first QTL includes a region between clusters of fatty acid desaturases (*Fads1*, *Fads2*, *Fads3*) and membrane-spanning 4A (*Ms4a*) gene members (**Fig. S2C**), which have literature support for associations to lipid metabolism, total cholesterol, and HDL-C (30–32). The second QTL, on Chr 19 included two compelling genes (**Fig. S2D**); N-acylsphingosine amidohydrolase 2 (*Asah2*) and Apobec1 complementation factor (*A1cf*). Due to associations of ceramides with cardiovascular disease (33), we sought to investigate *Asah2,* which encodes a ceramide-catabolizing neutral ceramidase, as a causal gene for HDL-2b lipoproteins.

### Validation of *Asah2* as a novel driver of plasma lipoprotein classes

To validate *Asah2* as a driver of HDL lipoproteins, whole-body *Asah2* knockout (*Asah2^-/-^*), heterozygous (*Asah2^+/-^*), or wild-type (*Asah2^+/+^*) mice were fed a high-fat diet (HFD) for 16 weeks and plasma collected at 25 weeks of age. Loss of *Asah2* gene expression was confirmed in liver tissues of mice from each sex and genotype (**Fig. S3**). To assess potential shifts in lipoprotein classes, we used ion mobility analysis to measure lipoprotein concentrations by size. Overall, female *Asah2^-/-^* mice had increased levels of all lipoproteins compared to female *Asah2^+/+^*mice, with significant increases in HDL and IDL sub-classes (**Fig. 4, Fig. S4,** p<0.01). Specifically, female *Asah2^-/-^* had significantly higher concentrations of small HDL-3,2a (**Fig. 4A**, p<0.009) and strong trends for large HDL-2b (**Fig. 4B**, p<0.07) particles. Both male and female *Asah2^-/-^* mice showed trends for increased particles in the midzone size range compared to *Asah2^+/+^*mice (p<0.08, p<0.09, respectively). Three LDL subclasses, LDL-IIB (**Fig. 4D**), LDL-IIa (**Fig. 4E**) and LDL-1 (**Fig. 4F**), were increased in female *Asah2^-/-^* versus *Asah2^+/+^*mice. Similarly, intermediate density lipoproteins, IDL-2 (**Fig. 4G**) and IDL-1 (**Fig. 4H**) were more abundant in female *Asah2^-/-^* mice (p<0.008). LDL-IIa, LDL-1 and both IDL particle classes were increased in female *Asah2^-/-^* versus *Asah2^+/-^* mice (p<0.04), highlighting a gene dosage effect for these particles. Finally, small VLDL (**Fig. 4I**) were also increased in female *Asah2^-/-^* compared to *Asah2^+/+^* mice (p<0.03). LDL, IDL and VLDL particle subclasses were not different between genotypes in male mice.

**Figure 4:**
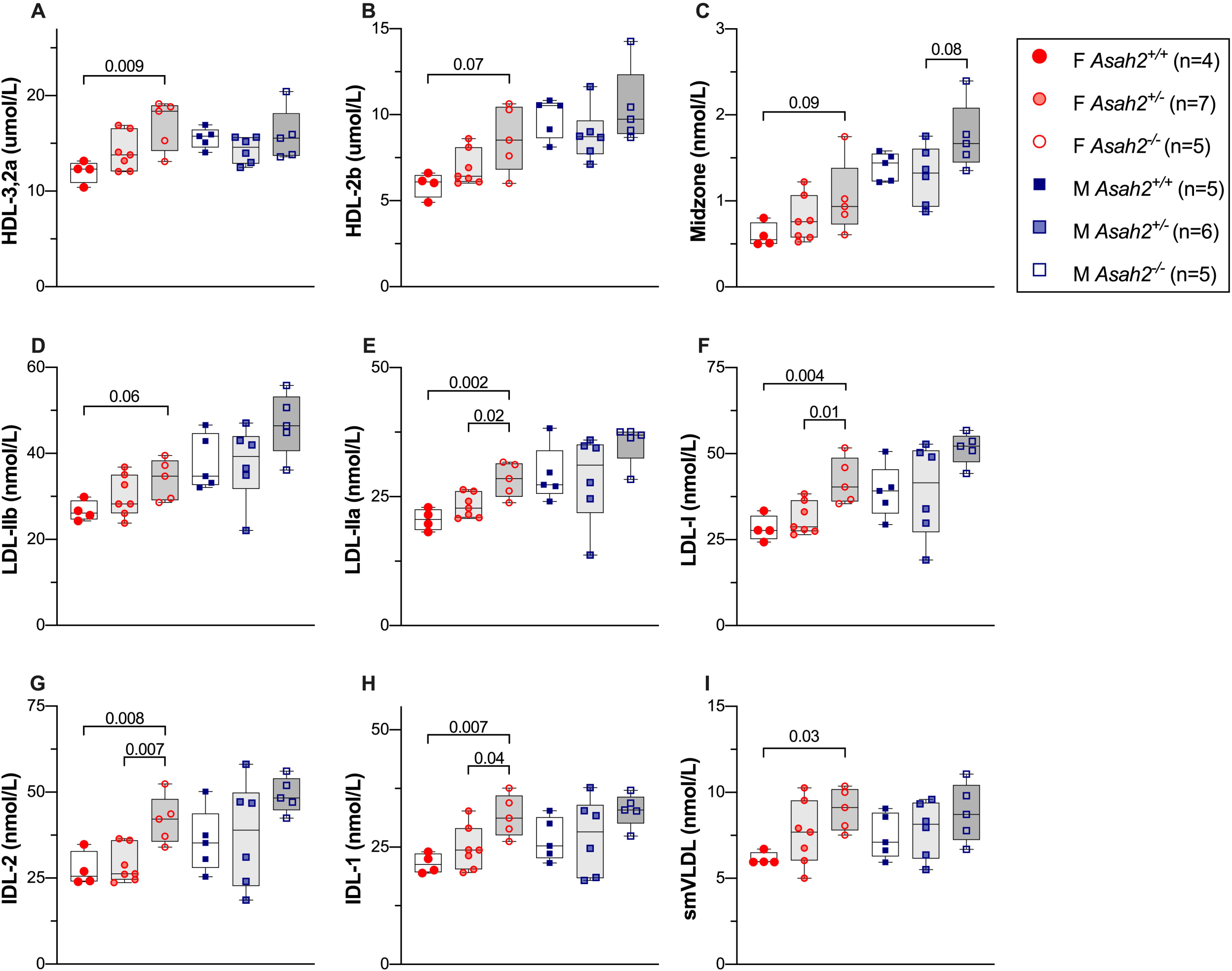
*Asah2* is a driver of plasma lipoproteins in female mice. Circulating lipoprotein subclasses measured by ion mobility analysis in female and male *Asah2^-/-^*, *Asah2^-/+,^*and *Asah2^+/+^* mice. (**A**) Small HDL-3,2a (76.5-105 Å) and (**B**) large HDL-2b (105-145 Å) concentrations were increased in *Asah2^-/-^*females. (**C**) Both male and female *Asah2^-/-^* showed a trend for increased particles in the midzone size range (145-180Å) compared to their *Asah2^+/+^* mice. Three LDL subclasses, LDL-IIB (**D)**, LDL-IIa (**E**) and LDL-1 (**F**), two intermediate density lipoproteins, IDL-2 (**G**) and IDL-1 (**H**), and small VLDL particles (**I**) were all elevated in *Asah2^-/-^* female mice.

Given the increased concentration of apo-B containing particles in *Asah2^-/-^* mice, we asked if the abundance of the LDL receptor (LDLR) protein or mRNA were altered in *Asah2^-/-^* mice. We assessed LDLR protein in liver by western blot. In *Asah2^-/-^* females, there was a small, but statistically significant increase in LDLR protein compared to *Asah2^+/+^* females (**Fig. S4F, S4H**); males were not different by genotype (**Fig. S4G, S4H**). Additionally, *Ldlr* and *Pcsk9* expression in liver was similar between genotypes of the same sex (**Fig. S4I**). Thus, the mechanism by which the loss of *Asah2* drives increased lipoprotein abundance is likely not due to downregulation of the LDLR protein.

### Integration of mouse lipoprotein QTL with human GWAS

To determine the translational significance of our findings to humans, we analyzed the syntenic loci for significant traits in human genome-wide associated studies (GWAS). To determine synteny, a 2-Mbp flanking region was first identified for each mouse lipoprotein QTL. This 2-Mbp region was then used with the LiftOver utility from UCSC Genomic Institute (genome.ucsc.edu/cgi-bin/hgLiftOver) to yield the human syntenic locus for each QTL (**Table S4**). We then asked if these loci were associated with cardiometabolic phenotypes, including cardiovascular, glycemic, blood lipid, or anthropometric (excluding height) traits in GWAS Central (www.gwascentral.org) (34).

Approximately 2,000 SNPs with cardiometabolic traits were identified within the syntenic regions (**Table S5**). SNPs syntenic with 20 of the 21 lipoprotein QTL identified in mice were strongly associated with metabolic traits in human GWAS (p<10^-8^, **Fig. 5**). Fifteen of the 21 loci were highly enriched with associations for three or more trait categories surveyed, providing evidence of significant genetic associations in the human population.

**Figure 5:**
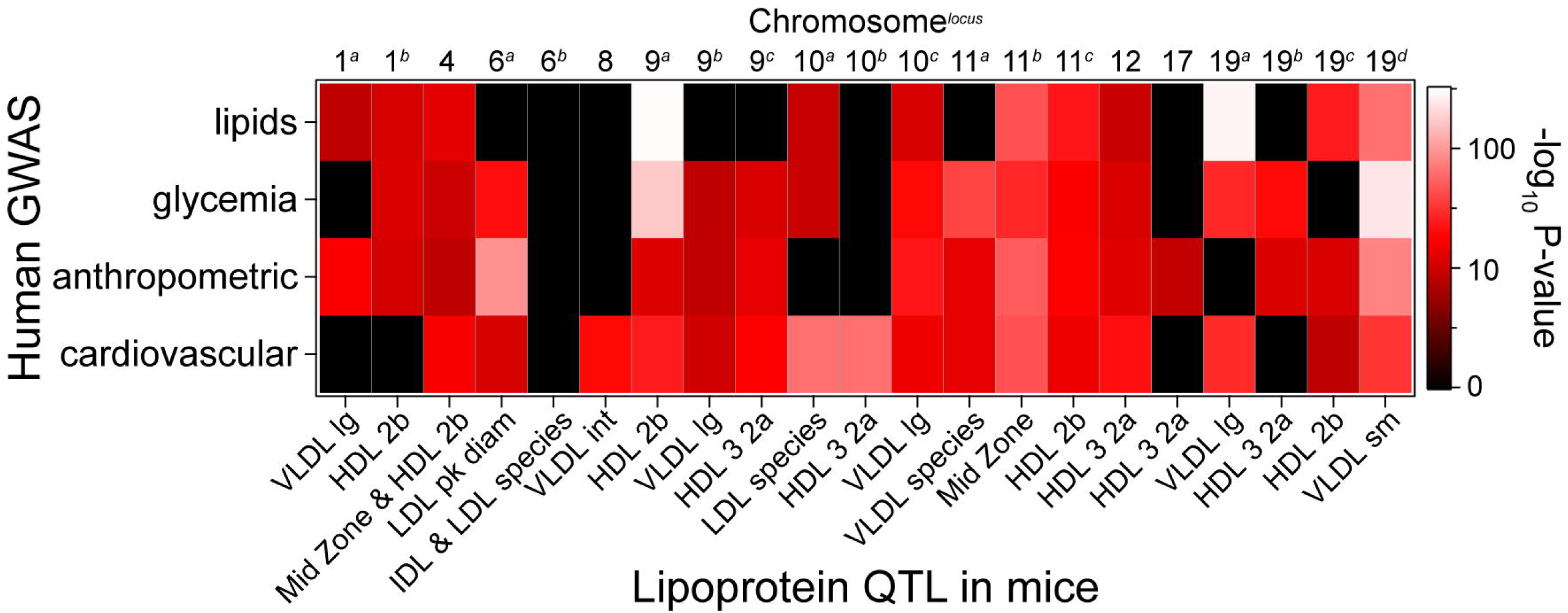
Mouse lipoprotein QTL are syntenic to regions associated with cardiometabolic traits in humans. Heatmap illustrates maximum enrichment (-log_10_ p-value) for SNPs that are present within regions syntenic to mouse lipoprotein QTL and associated with lipid, glycemia, anthropometric (excluding height) or cardiovascular phenotypes in human GWAS.

To nominate causal genes for each locus, we considered lipid-associated SNPs in mouse and human data and searched for literature support and mouse phenotyping data through the International Mouse Phenotyping Consortium (IMPC, www.mousephenotype.org). At mouse QTL, we prioritized SNPs that fell within the genomic region with a 1.5 LOD drop. At the syntenic loci in human, we identified SNPs with significant associations (p<10^-8^). Next, we searched published literature for phenotypic or mechanistic support. Through this integrated pipeline of mouse lipoprotein QTL, human GWAS data, and published, publicly available data, we nominated candidate driver genes for 18 of the 21 lipoprotein QTL (**Table 1**).

**Table 1:**
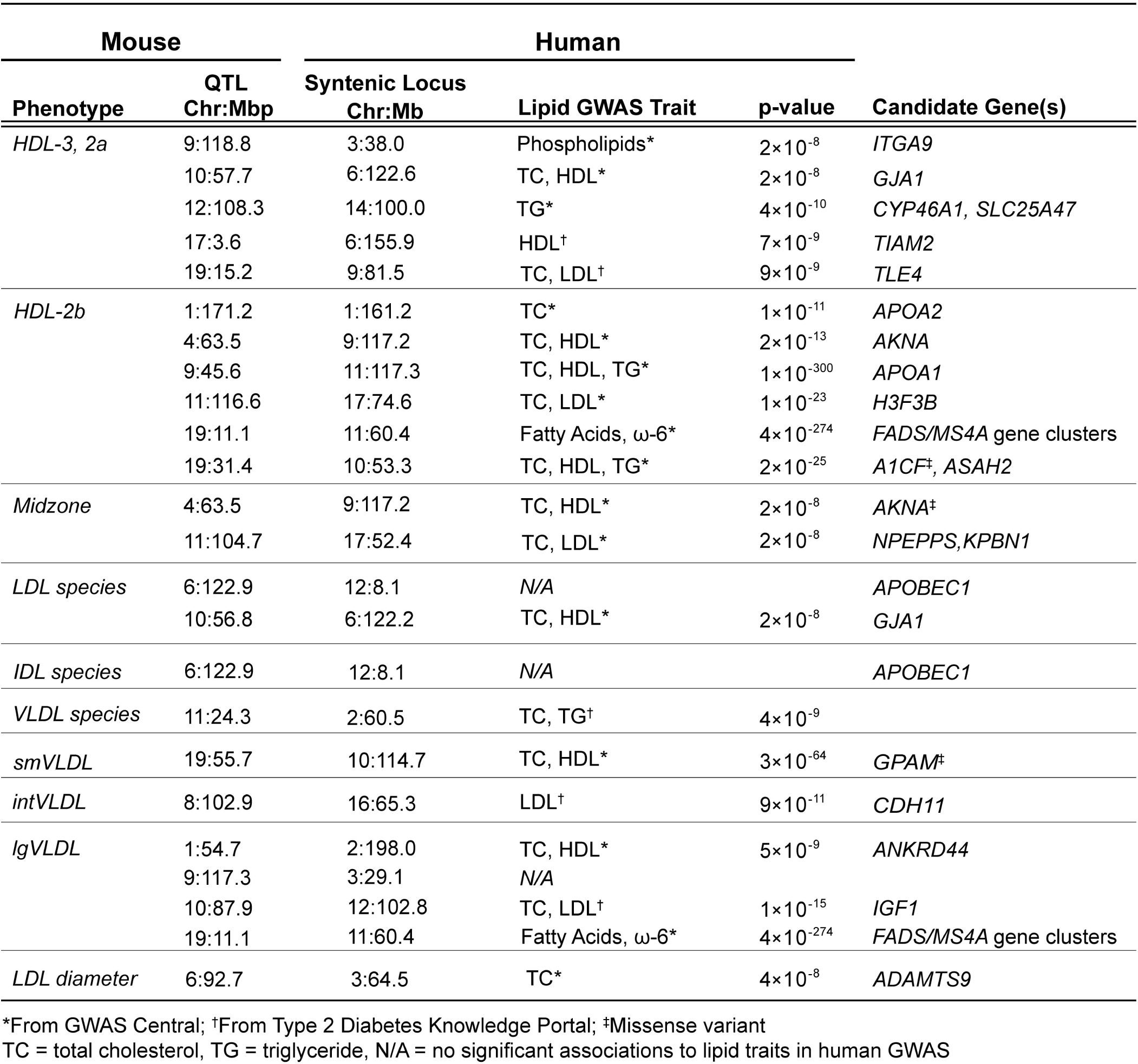
Candidate genes for plasma lipoprotein QTL. *From GWAS Central; †From Type 2 Diabetes Knowledge Portal; ‡Missense variant TC = total cholesterol, TG = triglyceride, N/A = no significant associations to lipid traits in human GWAS

| Mouse |  | Human |  |  |  |
| --- | --- | --- | --- | --- | --- |
| Phenotype | QTL | Syntenic Locus |  | p-value | Candidate Gene(s) |
|  | Chr:Mbp | Chr:Mb | Lipid GWAS Trait |  |  |
| *HDL-3, 2a* | 9:118.8 | 3:38.0 | Phospholipids* | 2×10<sup>-8</sup> | *ITGA9* |
|  | 10:57.7 | 6:122.6 | TC, HDL* | 2×10<sup>-8</sup> | *GJA1* |
|  | 12:108.3 | 14:100.0 | TG* | 4×10<sup>-10</sup> | *CYP46A1, SLC25A47* |
|  | 17:3.6 | 6:155.9 | HDL<sup>†</sup> | 7×10<sup>-9</sup> | *TIAM2* |
|  | 19:15.2 | 9:81.5 | TC, LDL<sup>†</sup> | 9×10<sup>-9</sup> | *TLE4* |
| *HDL-2b* | 1:171.2 | 1:161.2 | TC* | 1×10<sup>-11</sup> | *APOA2* |
|  | 4:63.5 | 9:117.2 | TC, HDL* | 2×10<sup>-13</sup> | *AKNA* |
|  | 9:45.6 | 11:117.3 | TC, HDL, TG* | 1×10<sup>-300</sup> | *APOA1* |
|  | 11:116.6 | 17:74.6 | TC, LDL* | 1×10<sup>-23</sup> | *H3F3B* |
|  | 19:11.1 | 11:60.4 | Fatty Acids, ω-6* | 4×10<sup>-274</sup> | *FADS/MS4A* gene clusters |
|  | 19:31.4 | 10:53.3 | TC, HDL, TG* | 2×10<sup>-25</sup> | *A1CF<sup>‡</sup>, ASAH2* |
| *Midzone* | 4:63.5 | 9:117.2 | TC, HDL* | 2×10<sup>-8</sup> | *AKNA<sup>‡</sup>* |
|  | 11:104.7 | 17:52.4 | TC, LDL* | 2×10<sup>-8</sup> | *NPEPPS, KPBN1* |
| *LDL species* | 6:122.9 | 12:8.1 | N/A |  | *APOBEC1* |
|  | 10:56.8 | 6:122.2 | TC, HDL* | 2×10<sup>-8</sup> | *GJA1* |
| *IDL species* | 6:122.9 | 12:8.1 | N/A |  | *APOBEC1* |
| *VLDL species* | 11:24.3 | 2:60.5 | TC, TG<sup>†</sup> | 4×10<sup>-9</sup> |  |
| *smVLDL* | 19:55.7 | 10:114.7 | TC, HDL* | 3×10<sup>-64</sup> | *GPAM<sup>‡</sup>* |
| *intVLDL* | 8:102.9 | 16:65.3 | LDL<sup>†</sup> | 9×10<sup>-11</sup> | *CDH11* |
| *lgVLDL* | 1:54.7 | 2:198.0 | TC, HDL* | 5×10<sup>-9</sup> | *ANKRD44* |
|  | 9:117.3 | 3:29.1 | N/A |  |  |
|  | 10:87.9 | 12:102.8 | TC, LDL<sup>†</sup> | 1×10<sup>-15</sup> | *IGF1* |
|  | 19:11.1 | 11:60.4 | Fatty Acids, ω-6* | 4×10<sup>-274</sup> | *FADS/MS4A* gene clusters |
| *LDL diameter* | 6:92.7 | 3:64.5 | TC* | 4×10<sup>-8</sup> | *ADAMTS9* |

We identified *Akna* as the candidate gene for HDL-2b and Midzone lipoprotein classes at a QTL on Chr 4. *Akna*, an AT-hook transcription factor (35), promotes *Cd40* expression (36), indicating a role in inflammatory processes. In human genetic association studies, *AKNA* variants have been linked to total cholesterol (37), HDL cholesterol and ApoA1 (38), and plasma sphingolipids (39). However, no studies have mechanistically linked *Akna* to cholesterol or lipoprotein metabolism. It is possible that *Akna* may reflect HDL’s role in inflammation and/or immune surveillance.

The second candidate driver for the HDL-2b locus on Chr 19 at ∼32 Mbp is *A1cf* (Apobec1 complementation factor). *A1cf* works in conjunction with *Apobec1* to catalyze editing of *Apob* mRNA, which results in the introduction of a stop codon and production of a truncated protein product, ApoB48 (40, 41). In human GWAS, there are strong associations between *A1CF* missense variants and plasma lipoprotein phenotypes (total cholesterol, LDL-C, serum ApoB) (34, 42).

We identified a single locus associated with both IDL and LDL, mapping to Chr 6 (**Figs. 1** and **2**). Interestingly, when we looked at the syntenic locus in humans, there were no significant associations for cardiovascular, lipid, anthropometric or glycemic traits. However, both the mouse and human genomic region contain *Apobec1* (apolipoprotein B mRNA editing enzyme catalytic subunit 1). In *Apobec1^-/-^* mice, plasma ApoB-100 is increased 176%, although this was not accompanied by changes to cholesterol concentrations in VLDL or LDL lipoprotein classes (43). The lack of genetic association in humans may be due to species differences where mice express *Apobec1* in both liver and intestine (44), while human expression is restricted to the intestine (45). It may also highlight the minimal contribution of plasma ApoB-48 concentration to total ApoB and the poor correlation to plasma cholesterol in humans (46, 47).

We mapped a single locus spanning five LDL subclasses (Chr 10) and one for LDL diameter (Chr 6). *Gja1,* also known as connexin 43 (*Cnx43*), is our candidate gene for the LDL species mapping to Chr 10*. Cnx43^+/-^*/*Ldlr^-/-^* mice have a reduction in atherosclerotic lesion formation, compared to *Cnx43^+/+^*/*Ldlr^-/-^*mice, without a change in plasma cholesterol or triglyceride (48). *Adamts9,* the candidate gene for LDL diameter, is a metallopeptidase with a type 1 thrombospondin motif. *Adamts9* has not been directly studied for a role in lipoprotein metabolism, but has been shown to be a suppressor of mTOR pathway in cancer cell lines (49). Both thrombospondin-1 and thrombospondin-2, however, have been studied in cardiovascular disease (50, 51) and mediate atherosclerotic plaque development in mice (52, 53).

We identified seven QTL for the 3 major size ranges within VLDL particles. At these loci, we identified five candidate genes. Small VLDL, which mapped to Chr 19, includes the gene *Gpam* at the human syntenic locus. In mice, the founder strains CAST and PWK have splice and untranslated region variants (Table S5), consistent with those two strains carrying the low alleles for the Chr 19 QTL (Table S1). Within human GWAS, *GPAM* is strongly associated with HDL, LDL, and total cholesterol, with missense variants that are highly associated to each of the three traits (Table S4, (42)). *Gpam* encodes for outer mitochondrial membrane glycerol-3-phosphate acyltransferase, an enzyme in the triglyceride synthesis pathway. Consistent with a role in lipoprotein metabolism, *Gpam* knockout mice have lower VLDL secretion rates and decreased liver triglycerides relative to wild-type mice (54).

We nominate *Igf1* as the causal gene at a QTL on Chr 10 for large VLDL. In mice, *Igf1* was shown to reduce liver cholesterol accumulation by activating *Abca1* (*55*) and an association between plasma *Igf1* cardiovascular disease risk has been found in human subjects (56). In bovine hepatocytes, *Apob* expression and VLDL secretion are increased with addition of exogenous Igf1 (57).

For two additional VLDL QTL (Chrs 1 and 8), we identified two lesser-known gene candidates: *Ankrd44* and *Cdh11*, respectively. In a study looking for genetic drivers of stroke in cerebral arteries, *Ankrd44* was down-regulated in veins from rabbits with hypercholesterolemia alone and with combined hypercholesterolemia and hypertension (58). *ANKRD44* genetic variants are associated with stroke risk in African Americans (59), though no mechanistic studies were found. Cadherin-11 (*Cdh11*), a cell adhesion protein, was nominated as the causal gene for intermediate-sized VLDL mapping to Chr 8. *Cdh11* has been implicated in autoimmune disorders, aortic valve calcification, and recently, scarring following myocardial infarction through its effect on fibrosis and inflammation(60). In a mouse model of atherosclerosis, *Cdh11* expression was increased in atherosclerotic plaques of *ApoE*^-/-^ mice. *ApoE*^-/-^/*Cdh11*^-/-^ double knockout mice had altered immune cell populations and increased atherosclerosis (61).

## Discussion

Lipoproteins that are isolated based on size or buoyant density represent snapshots of dynamic processes involving the transfer of lipids and proteins between particles and between tissues and lipoprotein particles. Genetic linkage and association studies help to identify the genes that affect these dynamic processes. The heterogeneity of the major lipoprotein classes (VLDL, LDL, HDL) in the DO founder strains suggested that we might find QTL that explain this heterogeneity (**Fig. 1**).

Prior to embarking on the genetic study in DO mice, we first estimated heritability (*h^2^*) of these distinct lipoprotein subclasses in the eight founder strains fed either chow or HFHS diets. Heritability estimates were reduced by approximately half for mice maintained on the HFHS diet (**Fig. 1C, D**), which in part may reflect increased intra-strain variance in particle concentrations (**Fig. S1**). Previous reports have highlighted an interaction between diet and genetics that alters heritability of metabolic phenotypes. In pedigreed baboons, high fat and/or high cholesterol diets increased variance and reduced *h^2^* in plasma HDL-C, median HDL size, and ApoA1/ApoB protein abundance compared to a low-fat, low-cholesterol diet (62). In a survey of 13 inbred mouse strains fed a Western diet, up to 75% of variance in lean body weight could be attributed to genetics (*h^2^*), whereas other phenotypes (e.g., plasma glucose) showed reduced heritability (*h^2^* < 20%) (63). Finally, a panel of 22 inbred CC strains showed that some traits (e.g., body weight and plasma cholesterol) showed higher variation than other traits (e.g., body fat) between high-protein or high-fat diets (64). Our results are consistent with these previous findings, showing that a metabolically challenging diet can increase non-genetic variance and thus reduce estimated *h^2^*of metabolic phenotypes.

Despite the reduction in estimated heritability observed for mice maintained on the HFHS diet, we identified 21 QTL for 16 lipoprotein subclasses in DO mice. Some of the loci contain genes encoding well-known apolipoproteins (*e.g.*, the locus harboring apoA1, C3, A4, A5, and PCSK7, and another locus containing apoA2), inspiring confidence in the ability of our screen to detect relevant loci. Nearly all these QTL, when lifted over into the human genome, are associated with lipid traits, as well as other cardiometabolic phenotypes, including cardiovascular disease risk, body weight, and diabetes.

At the HDL-2b locus on chromosome 19 at ∼32 Mbp, we identified two candidate drivers: *Asah2* and *A1cf.* We chose to investigate the relationship between *Asah2* and lipoproteins by phenotyping *Asah2^-/-^* mice. In female mice fed a high-fat diet, *Asah2* deletion resulted in increased HDL, midzone, large LDL, IDL, and small VLDL particles (**Fig. 4**). A similar trend was seen in midzone and IDL-2 particles in male mice. *Asah2* is highly expressed in the intestine and functions as a neutral ceramidase. Although sphingolipids, including ceramides, can be carried on LDL and VLDL particles (65), to our knowledge, this is the first time a ceramidase has been shown to affect lipoprotein abundance.

Ceramides have recently been recognized as a cholesterol-independent lipid biomarker of cardiovascular disease risk (66), with the potential for being a predictor of endothelial dysfunction and early atherosclerosis (67). In a mouse model of atherosclerosis, pharmacological inhibition of HIF-1α in adipose tissue decreased ceramide formation through a neutral sphingomyelinase and was associated with decreased plasma cholesterol and delayed atherosclerotic plaque progression (68). Moreover, targeted disruption of ceramide synthesis similarly blunts plaque formation. Specifically, genetic deletion(69) or pharmacological inhibition (70, 71) of serine palmitoyltransferase, which catalyzes the committed step in *de novo* ceramide synthesis, also blunts plaque formation. *Asah2* is required for intestinal degradation of dietary sphingolipids (19). As ASAH2 expression is predominantly in the intestine, its activity may alter intestinal absorption of cholesterol. Indeed, strategies that reduce ceramides in the gut, can blunt cholesterol absorption (72, 73). Moreover, ASAH2 mediates the body’s response to microbial sphingolipids (19, 74). Deletion of *Asah2* alleviates diet-induced NASH/NAFLD via down-regulation of stearoyl-CoA desaturase (*Scd1*) and reduces cholesterol accumulation (75). When fed a very low fat diet, *Scd1^-/-^* mice have increased plasma cholesterol, especially in the LDL and VLDL fractions (76). When fed a standard chow diet (13% calories as fat), *Scd1^-/-^* mice have increased plasma HDL-C (77). Therefore, one might hypothesize that *Asah2* modulates lipoprotein metabolism through its effect on *Scd1* expression.

For many years, elevated HDL was considered to be protective against atherosclerosis. However, Mendelian randomization studies and mouse knockout experiments showed this concept was overly simplistic (78). Rather, opposing dynamic processes affect HDL and atherosclerosis risk by affecting the dynamics of cholesterol transport. For example, mutations in *ABCA1* reduce the ability of cells to transport cholesterol and phospholipids out of cells (79, 80). This results in lower HDL and increased atherosclerosis (81). In contrast, mutations in *SRB1* decrease the transport of cholesterol esters from HDL into cells (82–84). This leads to increased HDL and increased atherosclerosis.

We are not aware of any genetic association between *ASAH2* and atherosclerosis. Its modulation of HDL could increase or decrease atherosclerosis. Alternatively, *ASAH2* could affect atherosclerosis through its effects on sphingolipids, independent of its effect on HDL, through their effects on inflammatory pathways (85). Indeed, ceramides predict coronary artery disease independently of cholesterol (86). Pharmacological inhibition of ceramide synthesis via the serine palmitoyltransferase inhibitor, myriocin, reduces atherosclerotic plaque formation in APOE-deficient mice (71, 73). Moreover, ceramidases can also increase the formation of sphingosine-1-phosphate, an anti-atherogenic lipid which is largely carried in ApoM-containing particles (87).

In summary, we mapped mouse lipoprotein subclasses to identify 21 unique QTL which we then cross-referenced with the human syntenic locus to determine candidate genes associated with cardiometabolic traits in human GWAS. By integrating mouse data with human GWAS at the syntenic locus, we nominated candidate genes for 18 unique lipoprotein QTL. Deletion of *Asah2,* a novel candidate driver for HDL-2b, resulted in increased HDL, midzone, large LDL, IDL, and small VLDL particles in plasma from female mice. Similar validation experiments can be performed to explore the candidate genes we have nominated at the remaining QTL.

## Data Availability

All data for this study are provided in the main text or as supporting information.

## Acknowledgements

We appreciate the gift of *Asah2^-/-^*mice by Richard Proia (NIH) to the Summers/Holland lab.

## Funding

This work was supported by grants from the NIH (R01DK101573, R01DK102948, and RC2DK125961 (A.D.A.)) and by the University of Wisconsin–Madison, Department of Biochemistry and Office of the Vice Chancellor for Research and Graduate Education with funding from the Wisconsin Alumni Research Foundation (M.P.K.). Research support to T.R.P. was provided through the NIH by the Training Program in Translational Cardiovascular Science (T32-HL007936) at UW-Madison. Support to C.H.E. was provided by the American Diabetes Association (7-21-PDF-157).

## Abbreviations

CC: Collaborative Cross
Chr: chromosome
DO: Diversity Outbred
eQTL: expression quantitative trait locus
FPLC: fast protein liquid chromatography
GWAS: genome wide association studies
HFD: high-fat diet
HFHS: high-fat, high-sucrose diet
IMA: ion mobility spectrometry analysis
IMPC: International Mouse Phenotyping Consortium
LDLR: low-density lipoprotein receptor
LOD: logarithm of odds
Mbp: megabase pair
Pcsk7: Proprotein convertase subtilisin–kexin type 7
QTL: quantitative trait locus
TG: triglyceride

---

**Figure S1: Genetics and diet converge to influence circulating lipoproteins.**

Circulating concentrations (pM) for 16 lipoprotein subclasses and LDL peak diameter (Å) are illustrated for both sexes of the eight DO founder strains maintained on either regular rodent chow diet, or a Western-style diet high in fat and sucrose. Subclasses are ordered from small (**A**) to large (**B**) sizes.

**Figure S2: SNP association profiles for four HDL-2b QTL**

SNP association profiles and nearby genes at four QTL for HDL-2b. Chr 1 with 31 genes, including *Apoa2* (**A**); Chr 9 with 48 genes, including *Apoa1*, *Apoa4* and *Apoc3* genes (**B**); and two QTL on Chr 9, with 40 genes (**C**) and 24 genes (**D**).

**Figure S3: Confirmation of *Asah2* deletion in liver**

Loss of *Asah2* mRNA was confirmed in liver tissue of wildtype (*Asah2^+/+^*), heterozygous (*Asah2^+/-^*), or whole-body knockout (*Asah2^-/-^)* female and male mice.

**Figure S4: Loss of *Asah2* increases lipoprotein concentrations in female mice.**

Summed concentration values for total HDL (**A**), Midzone (**B**), LDL (**C**), IDL (**D**), and VLDL (**E**) lipoproteins are illustrated for female and male wildtype (*Asah2^+/+^*), heterozygous (*Asah2^+/-^*), and whole-body knockout (*Asah2^-/-^)* mice maintained on a high-fat diet for 25 weeks. Size ranges for these lipoprotein classes are provided in **Table S1**. LDLR protein (**F – H**) and gene expression (**I**) were unchanged in liver tissue for genotypes of each sex.

## Notes

### Competing Interest Statement

The authors have declared no competing interest.

