## Supplementary figures and images for "Identification of genetic drivers of plasma lipoproteins in the Diversity Outbred mouse population"

### Supplemental Figure S1

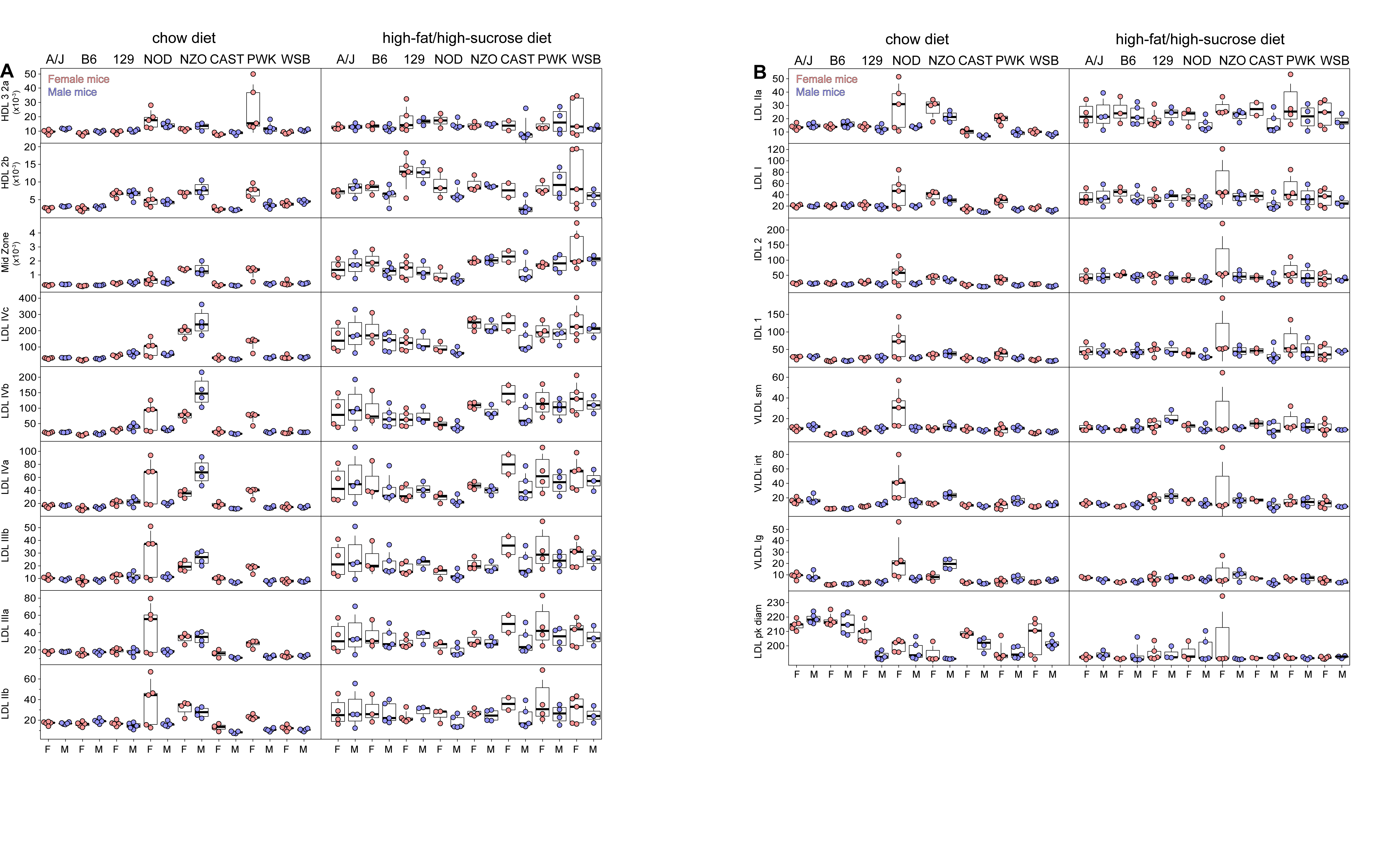

### Supplemental Figure S2

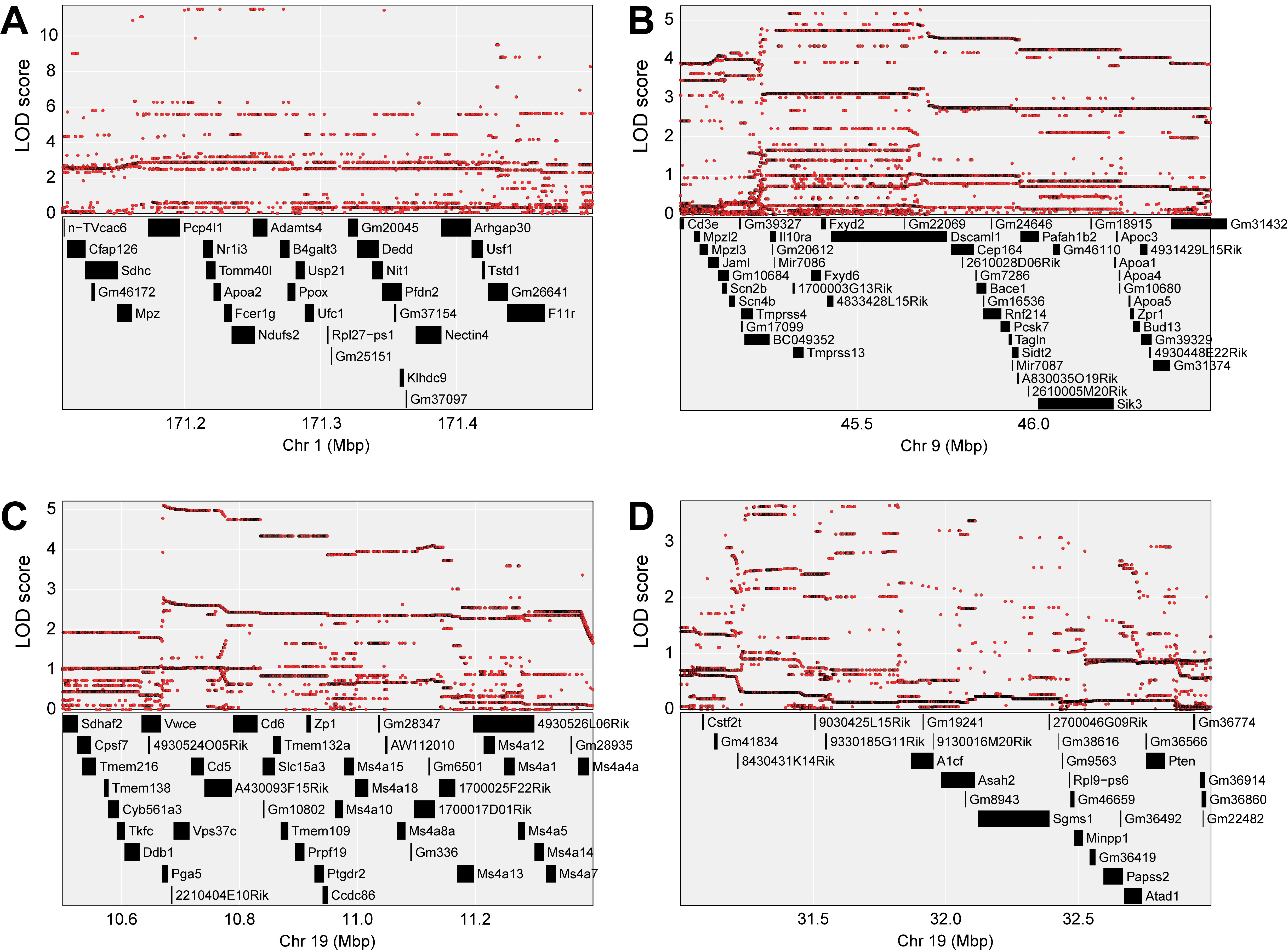

### Supplemental Figure S3

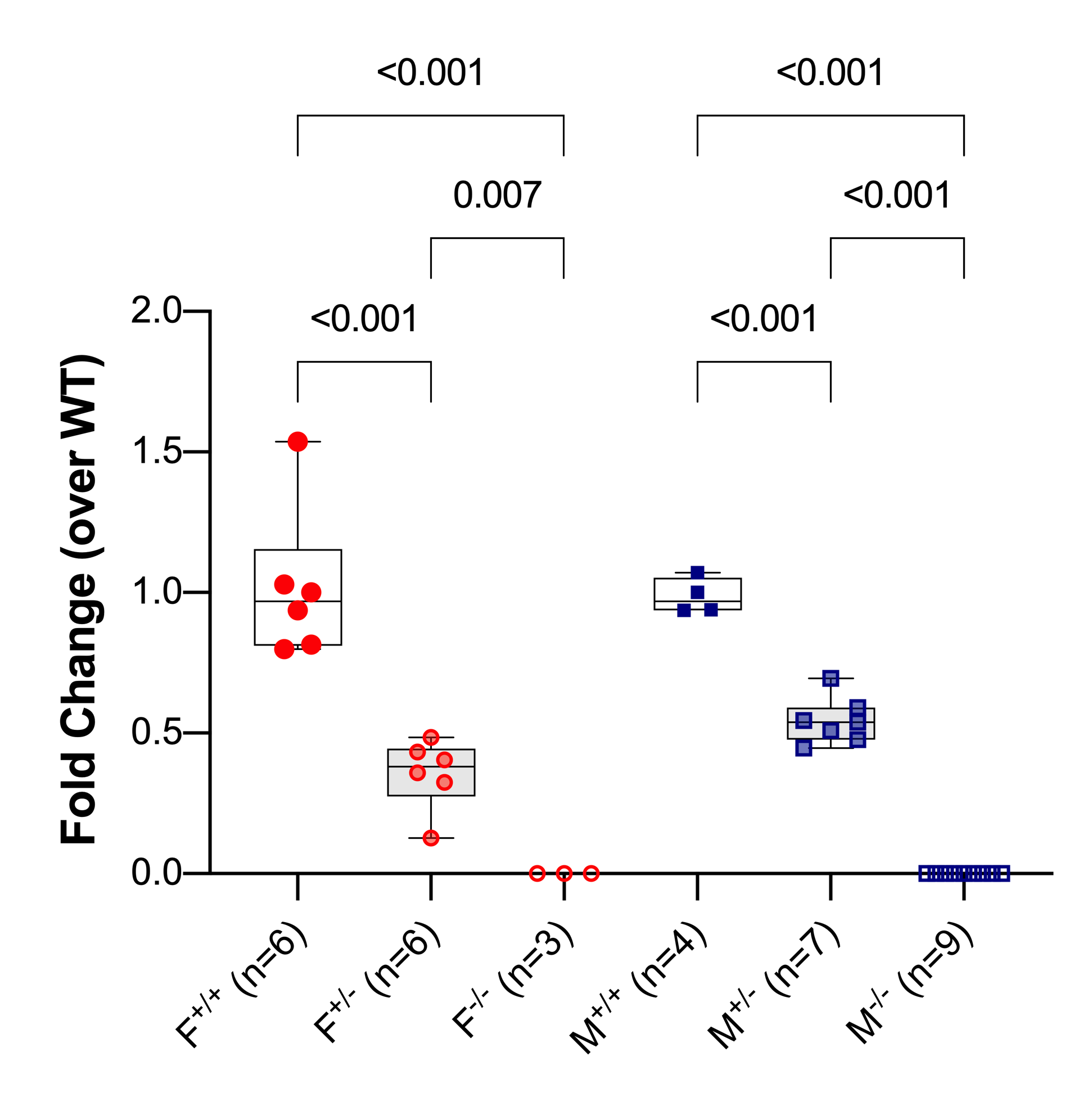

### Supplemental Figure S4

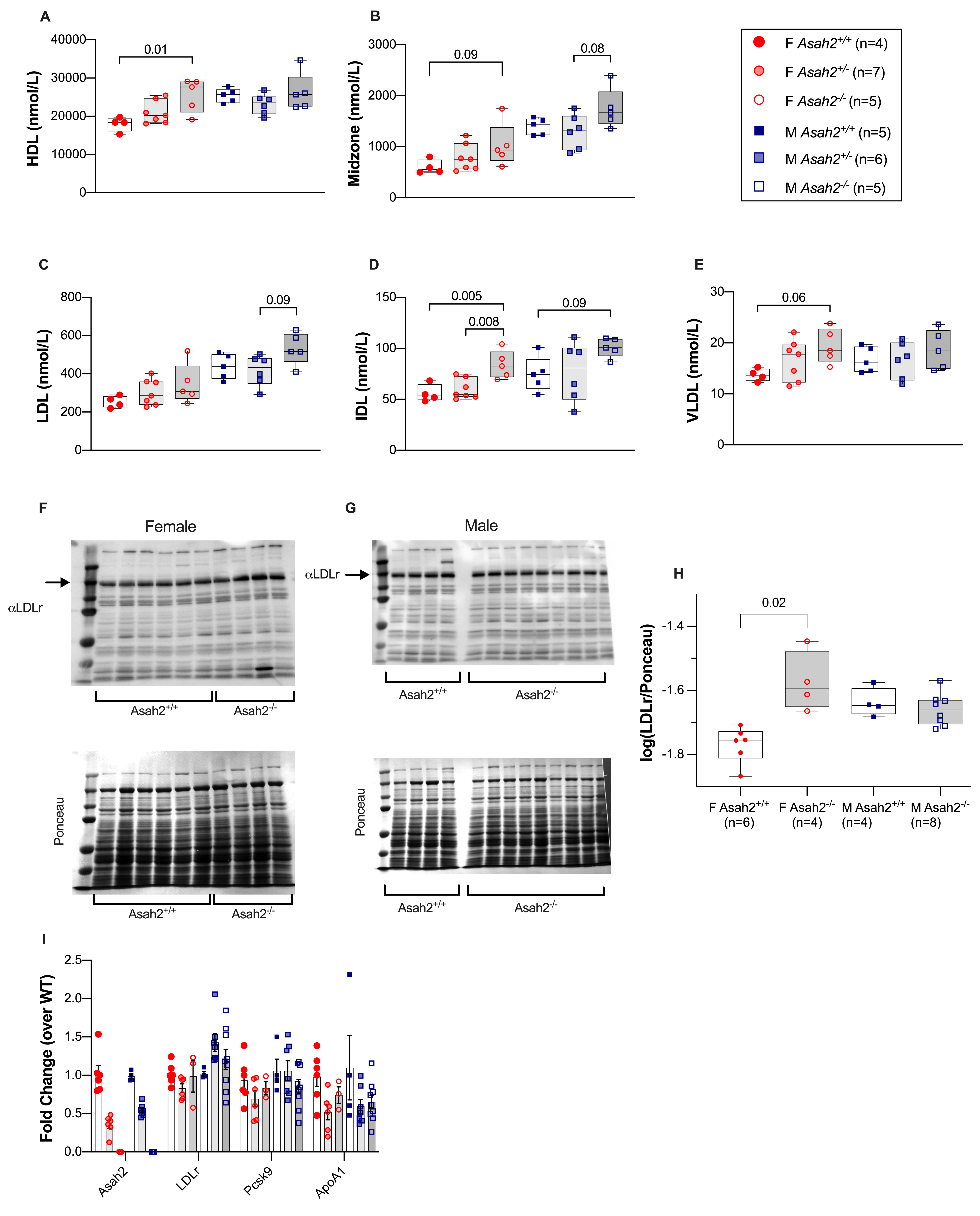
